# Experimental platform to study the neuronal mechanisms of magnetoreception in the honey bee

**DOI:** 10.64898/2026.05.05.722881

**Authors:** Alan Oesterle, Alara Kiriş, Albrecht Haase

## Abstract

Honey bees (*Apis mellifera*) offer an alternative model for investigating magnetoreception, exhibiting reliable navigation and behavioral responses to magnetic fields. Thanks to their compact brains, well-matched to the penetration depth of modern optical imaging techniques, they may offer insights into the neural and biochemical mechanisms underlying this sense, something that standard models, like migrating birds, have not so far provided. But also in honeybees, this progress requires tools capable of resolving weak, magnetically induced neural activity with high spatio-temporal precision. The approach, presented here, bridges quantum biology and neuroscience, allowing for testing the radical pair mechanism (RPM) as a potential basis for magnetic sensing. As the RPM predicts that magnetoreception is coupled to the visual system, we developed an *in vivo* two-photon calcium imaging approach to measure neural activity in the anterior optic tubercle, a higher-order visual center involved in chromatic processing and potentially navigation. Bees were prepared using a minimally invasive technique, in which this neuropil was retrogradely labelled with a fluorescent calcium indicator, enabling stable recording conditions over several hours. Controlled blue-light stimuli were provided by the scattered output of a fiber laser, and weak magnetic-field stimuli were applied by a shielded, three-axis Helmholtz coil system that allowed precise modulation of field strength and polarity while minimizing electromagnetic interference. Visual stimulation evoked consistent and reproducible calcium responses, validating the preparation and imaging stability. Magnetic stimulation produced small fluorescence decreases, suggesting field-dependent modulation of neural activity. The developed imaging framework shows the feasibility of detecting magnetic modulation in vision- and navigation-related brain regions, suggesting neural amplification of weak magnetic cues and providing a platform for controlled tests of RPM-specific predictions, including light dependence, polarity independence, and radiofrequency perturbation.

## 1 Introduction

To understand how animals detect and process the Earth’s magnetic field, one must take on the unique challenge of investigating the molecular and neural processes that link quantum-biological mechanisms to behavioral outputs [1]. Magnetoreception has been demonstrated across a wide range of taxa, including birds [2], rodents [3], sea turtles [4], and several invertebrates, such as fruit flies [5], ants [6], butterflies [7], cockroaches [8], and lobsters [9]. Despite this breadth of evidence, the mechanistic basis of magnetoreception and the associated neural pathways remain incompletely understood [10].

The honeybee (*Apis mellifera*) constitutes a particularly advantageous model organism for investigating these processes. Honeybees forage over distances exceeding 10 km [11] and rely on the integration of multiple navigational cues [12–14]. Behavioral evidence for magnetic field sensitivity has been reported in this species by multiple independent studies [15–21], and their well-characterized neuroanatomy [22] makes them well-suited for studying the neural correlates of magnetic sensing.

Three principal hypotheses have been proposed to explain magnetoreception [23,24]: magnetite-based mechanisms [25,26], electromagnetic induction [27], the radical pair mechanism (RPM) [28,29], or hybrid systems [30]. Although the RPM currently represents the most extensively investigated and widely supported model, evidence consistent with magnetite-based and induction-based mechanisms continues to emerge [31,32]. The RPM posits that magnetoreception arises from light-dependent photochemical reactions generating long-lived, spin-correlated radical pairs [23]. These radical pairs are sensitive to magnetic fields as weak as the Earth’s magnetic field (≈ 0.5 G), resulting in magnetically dependent changes in reaction yields that may influence downstream neural activity. Consistent with the observed light dependence of magnetoreception, it has been proposed that these reactions occur within the blue-light photoreceptor cryptochrome (CRY) [29,33–38]. Because cryptochrome is expressed in the retina, magnetic sensing has been hypothesized to interact with visual processing pathways, potentially manifesting as a modulation of visual input, perceived as a light-dark pattern superimposed onto the normal visual field [39,40].

To date, experimental investigations of the RPM have largely focused on behaviour [41,42], *in vitro* experiments [38,43], or heterologous expression of CRY outside its native cellular context [44]. In contrast, there are currently no tools available to directly measure magnetic-field-dependent neural activity in intact central brain circuits under controlled and physiologically relevant conditions. *In vivo* imaging approaches capable of resolving neuronal activity in central brain regions are predestined to address this gap [45–47].

Two-photon calcium imaging provides high spatiotemporal resolution and sufficient imaging depth to access neuronal activity in intact brain tissue [48]. The confined excitation volume reduces photobleaching and phototoxicity, while near-infrared excitation enables imaging at depths approaching 1 mm in scattering tissue [49]. Given that the honeybee brain has a volume of approximately 1 mm^3^ [22], this approach enables functional imaging across large portions of the brain, unlike in migrating birds, the standard model in magnetoreception research, whose larger brains limit optical access.

In this study, based on methods developed in previous studies [50–52], we establish an approach for *in vivo* two-photon calcium imaging of neuronal activity in the anterior optic tubercle (AOTu), a higher-order visual processing center involved in chromatic processing and navigation [53–57]. The AOTu receives input from the optic lobes via the anterior optic tract and is interconnected across hemispheres through the ventral intertubercular tract (vITT). Targeting this region may enable the investigation of magnetically mediated signals beyond the retina, providing access to neural circuits where signal integration and amplification occur naturally. This approach may open new avenues for research in magnetoreception, facilitating the exploration of downstream neural processing of weak primary signals, and providing a framework for examining how weak magnetic cues are transduced and integrated in the brain.

## 2 Methods

### 2.1 Bee Preparation

The honeybee preparation protocol was developed based on previous methods to image the bee’s visual system [55] and olfactory system [58].

#### 2.1.1 Neuronal staining protocol

1. Returning foragers were collected at the hive entrance between 10 am and 12 pm using a small plastic container. Bees were fed 5-6 µL of 50% (w/w) sucrose solution to maintain viability during preparation.
2. Bees were immobilized by chilling on ice until they stopped moving and were then mounted in a custom Plexiglass holder (Fig. 1). To stabilize the head, a wedge of soft dental wax (Zeta) was inserted between the head and the mount and shaped accordingly. The thorax was supported with a posterior sponge to limit body movement.
3. Antennae were restrained with thin, bent copper wires and secured at the base with n-eicosane (Thermo Fisher Scientific). The copper wires were then removed.
4. To expose the brain, a rectangular window was cut into a section of the head cuticle with a knife made from double-edged razor blades, which was secured in a razor blade holder (WPI).
5. The left salivary glands were displaced or removed, and tracheal segments adjacent to the left AOTu were carefully displaced to provide access for dye injection (Fig. 2A).
6. Crystallized Fura-dextran (10.000 MW, ThermoFisher Scientific) was loaded onto the tips of three borosilicate glass needles produced with a Pipette puller (Narishige) to a tip size of approximately 5 µm (Fig. 2B) and injected into the left AOTu using the mushroom body *α*-lobe as a landmark, as it has a slightly darker tone than the surrounding tissue (Fig. 2A).
7. The removed cuticle was replaced, and bees were fed an additional 3 µL of 50% (w/w) sucrose solution. Following injection, bees were maintained overnight in a dark, humid chamber to allow dye diffusion into the contralateral AOTu via the ventral intermediate tract (vITT).

**Fig. 1.**
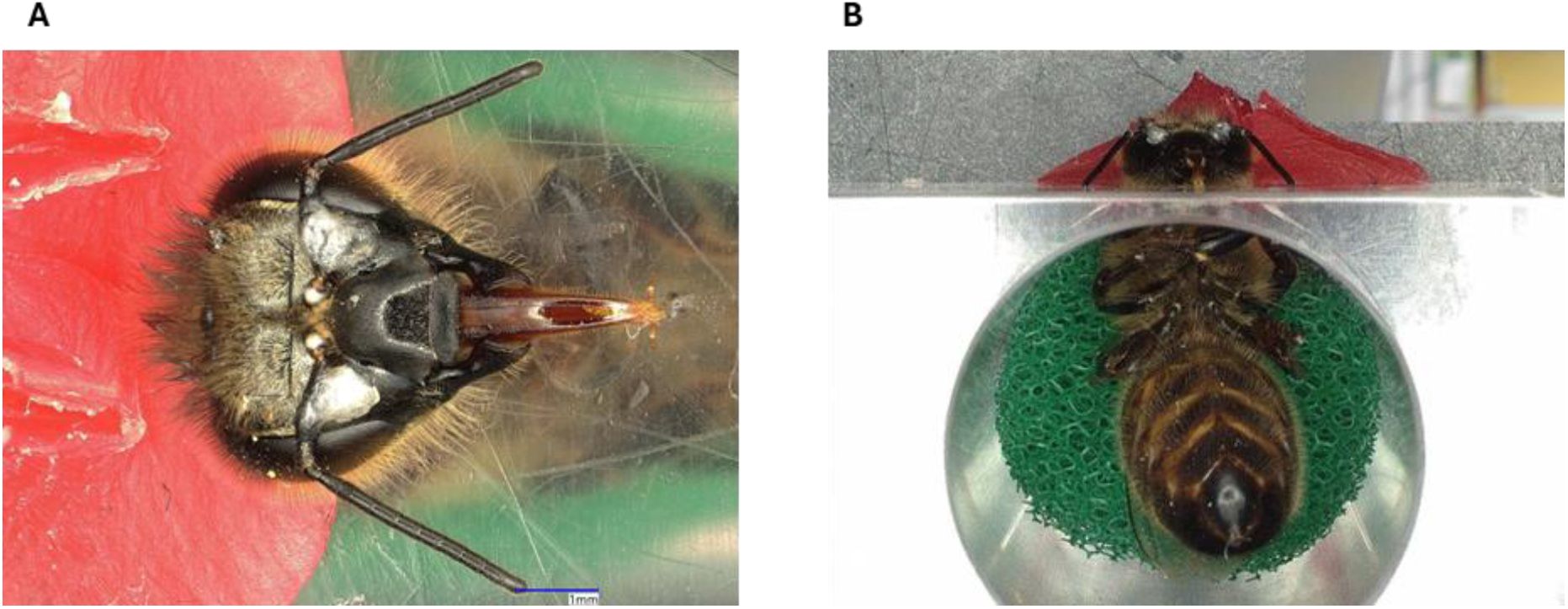
(A) Head fixation to the plexiglass mount with soft dental wax (red), (B) Thorax supported with a sponge (green).

**Fig. 2.**
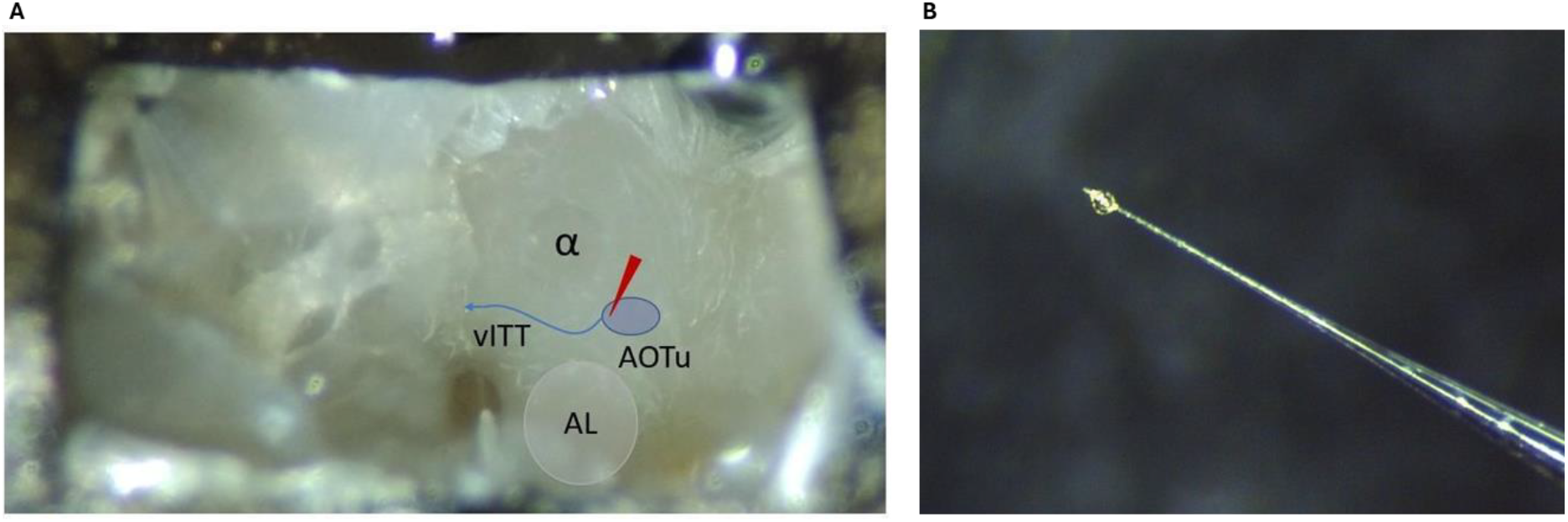
(A) Window in the cuticula exposing the bee brain, with tracheae cleared on the right side. The injection site is indicated by a red arrow. The bee’s antennae are facing downward. (B) Injection needle prepared with crystallized Fura at the tip.

#### 2.1.2 Imaging preparation protocol

1. The following morning, mandibles were immobilized with n-eicosane to minimize movement (Fig. 3). The head capsule was reopened, and the salivary glands and tracheae overlaying the right AOTu were carefully displaced.
2. Surface hemolymph was absorbed with tissue, and the head capsule was sealed with a droplet of transparent two-component silicone (Kwik-Sil, WPI). Before curing, a thin plastic sheet was placed in contact with the silicone, with edges bent upward, and coated with black acrylic paint, leaving a small observation window (Fig. 4A,B).
3. The bee was positioned under the objective of the two-photon microscope (Fig. 4C), with a water droplet placed between the water-immersion lens and the observation window.

**Fig. 3:**
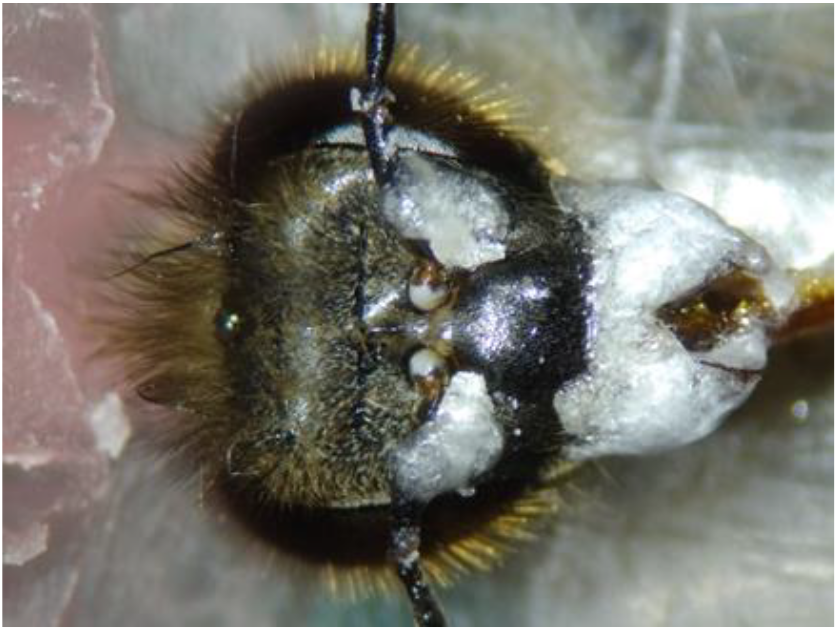
Bee head with antennae and mandibles immobilized using n-eicosane.

**Fig. 4:**
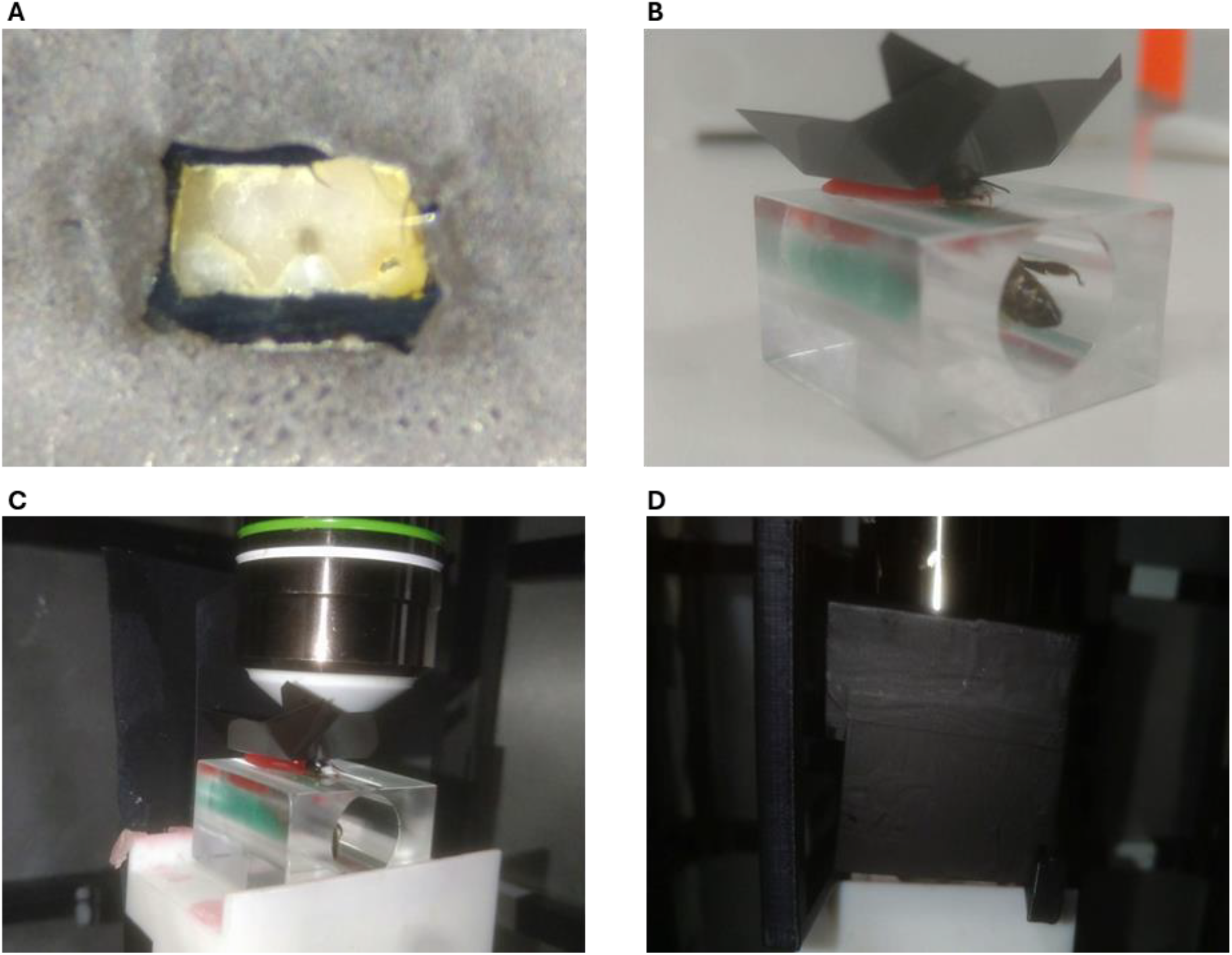
(A) Bee brain visible through a transparent window in the painted plastic shield, (B) Fully prepared bee with a black plastic shield against stray light, (C) Prepared bee, positioned under the microscope objective. (D) Full stray light shielding added.

This preparation enables stable recordings of neural activity for 3–4 hours, after which responses progressively diminish.

### 2.2 Visual stimulation setup

The visual stimulus was generated by a modulatable 473 nm diode laser (iBeam smart 473, TOPTICA) coupled to an optical fiber, providing up to 55 mW output power. The beam entered through an opening in the aluminum housing and was scattered by the rear panel, resulting in a diffuse, uniform illumination of the side panel, the visual field of the bee eye, which was enhanced by placing sheet of white paper on the surface (Fig. 5). The photon flux incident on the bee’s eye, measured with an amplified photodiode, was approximately 4 × 10^17^ photons m^−2^s^−1^, which corresponds to about 1 % of the intensity of the blue component in daylight. To minimize interference with fluorescence detection, the whole preparation was enclosed in a shielding structure made from rigid plastic foil painted black with acrylic paint (Fig. 4D). Two 473-488 nm notch filters (OD *>* 6.5, Chroma) were placed before the microscope’s photomultiplier tubes to remove residual stray light. The laser power for stimulation was usually about 20 mW. Each experimental session comprised 15 trials with a 3–5 s visual stimulus and a 26 s inter-trial interval.

**Fig. 5.**
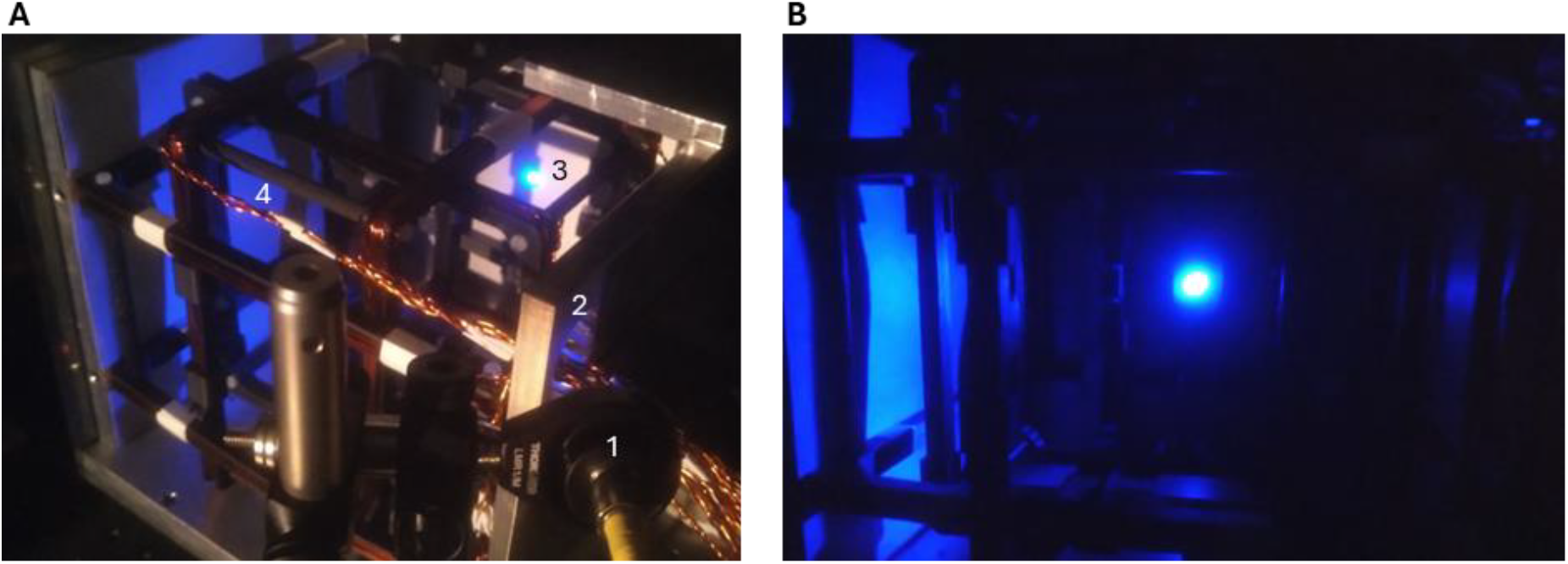
Visual stimulation setup: (A) side view showing the laser path from the fiber output (1) through the hole in the shielding cage (2), scattered from the rear panel (3), and producing a homogeneous illumination of the side panel (4). (B) Front view showing the stimulus under experimental conditions. The illuminated left side panel covers the field of view of the bee’s eye.

### 2.3 Magnetic stimulation setup

Magnetic fields were produced using a custom three-axis system of nested square Helmholtz coils (Fig. 6). The outermost coil pair was separated by 90 mm to accommodate nesting. The inner, middle, and outer coils had nominal widths of 154, 168, and 182 mm, with corresponding gaps of 75, 83, and 90 mm.

**Fig. 6.**
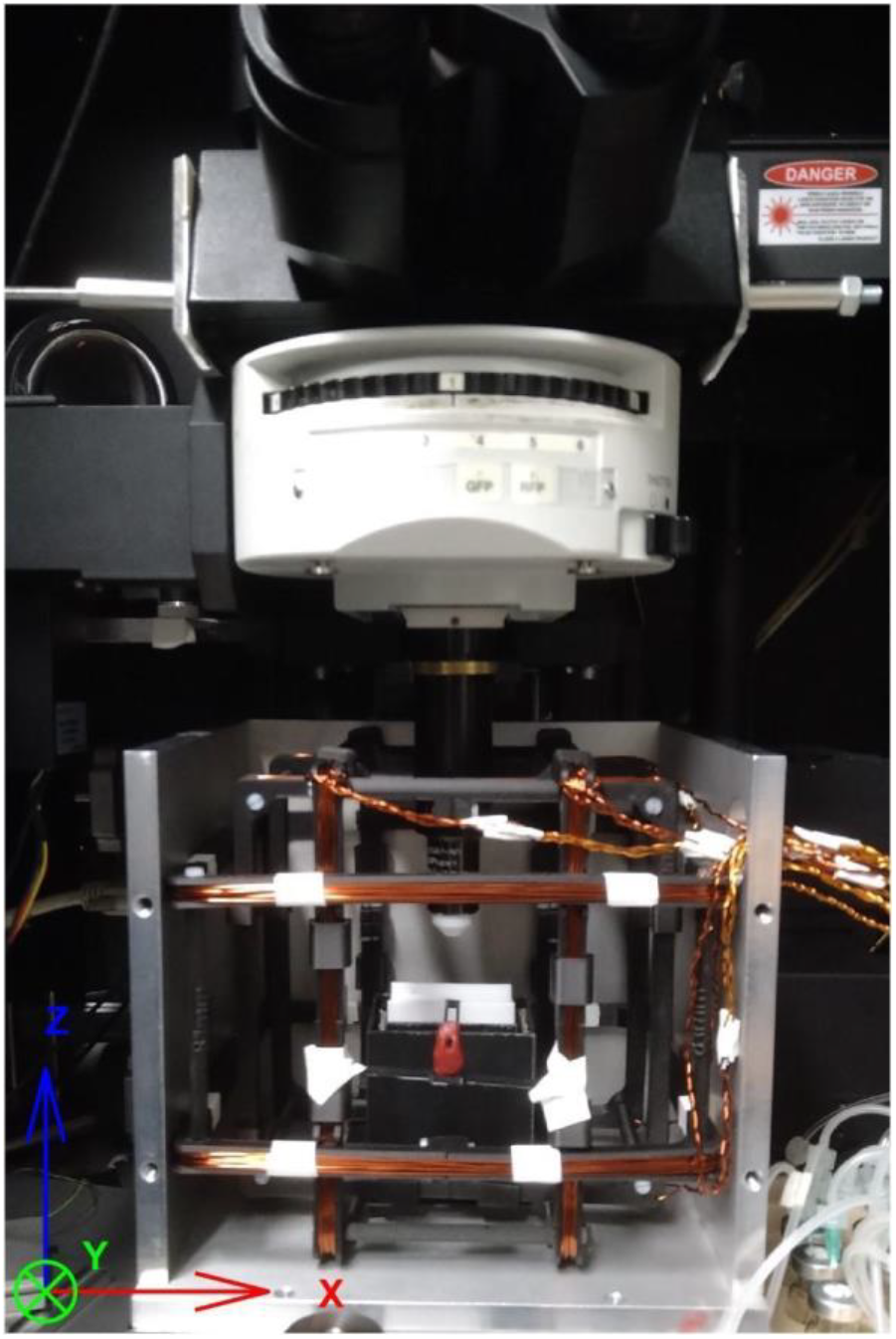
Two-photon microscope with the coil assembly and the shielding cage, with the upper and front plates removed. The magnetic field axes are labeled at the bottom left.

Field homogeneity was verified by simulation. Each coil comprised 26 windings of 1 mm copper wire distributed over five layers. Following Ahlers et al. [59], coils were double-wrapped, with two parallel windings per coil that could be electronically switched between series (field-adding) or series-opposition (field-cancelling) configurations. This allows for a sham stimulus to discriminate between genuine magnetic effects and electrical artifacts. A power supply (Rohde & Schwarz, NGE103) with remote control capability was used to generate fully customizable magnetic stimulus sequences. To reduce electromagnetic interference, the coils were enclosed in a 1 cm thick aluminum housing providing radio-frequency shielding, supplemented with 0.5 mm *µ*-metal sheets (Holland Shielding) on the exterior to attenuate static magnetic fields. The small openings in the shield allow for minimal residual ambient fields, which were compensated for by the coil system. Under zero-field conditions, the residual magnetic field could be reduced to 1.3 mG, 0.2 mG, and 0.9 mG along the X, Y, and Z axes, respectively, with standard deviations of 0.7, 0.6, and 0.7 mG (Fig. 7).

**Fig. 7.**
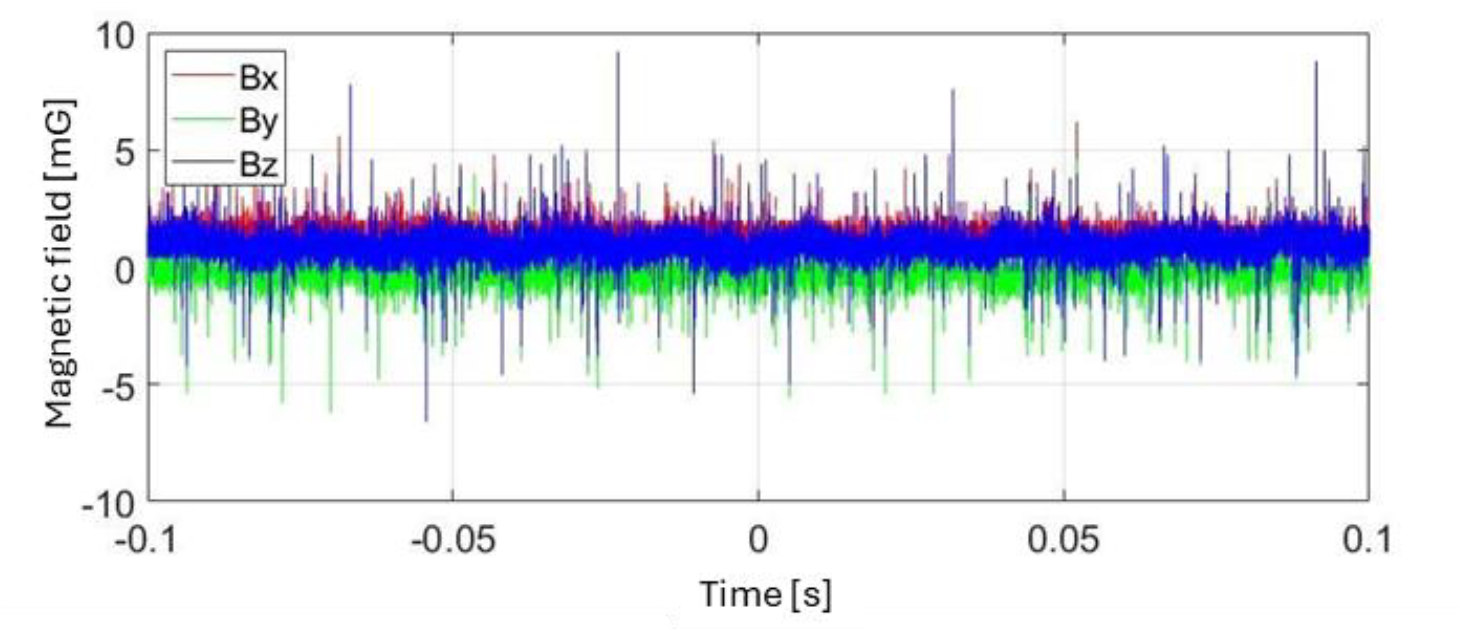
Measurement of fluctuations and offset under zero magnetic field conditions at the center of the Helmholtz coils.

Magnetic stimulation was generated, starting from a zero-field baseline, by applying a 0.5 G field along the X axis. To suppress high-frequency noise from abrupt current changes, the current was linearly ramped over 0.5 s (Fig. 8). The stimulus sequence for each trial involving magnetic stimulation was structured as follows: after an abituation time, first, the light was turned on for 13 s, followed by a 3 s magnetic stimulus, 10 s later also the light was switched off (Fig. 9). This sequence design allowed us from the the light ON/OFF responses to evaluate if the AOTu remained responsive throughout each trial. The total trial duration was 52 s, and 10 trials were recorded per stimulation condition.

**Fig. 8.**
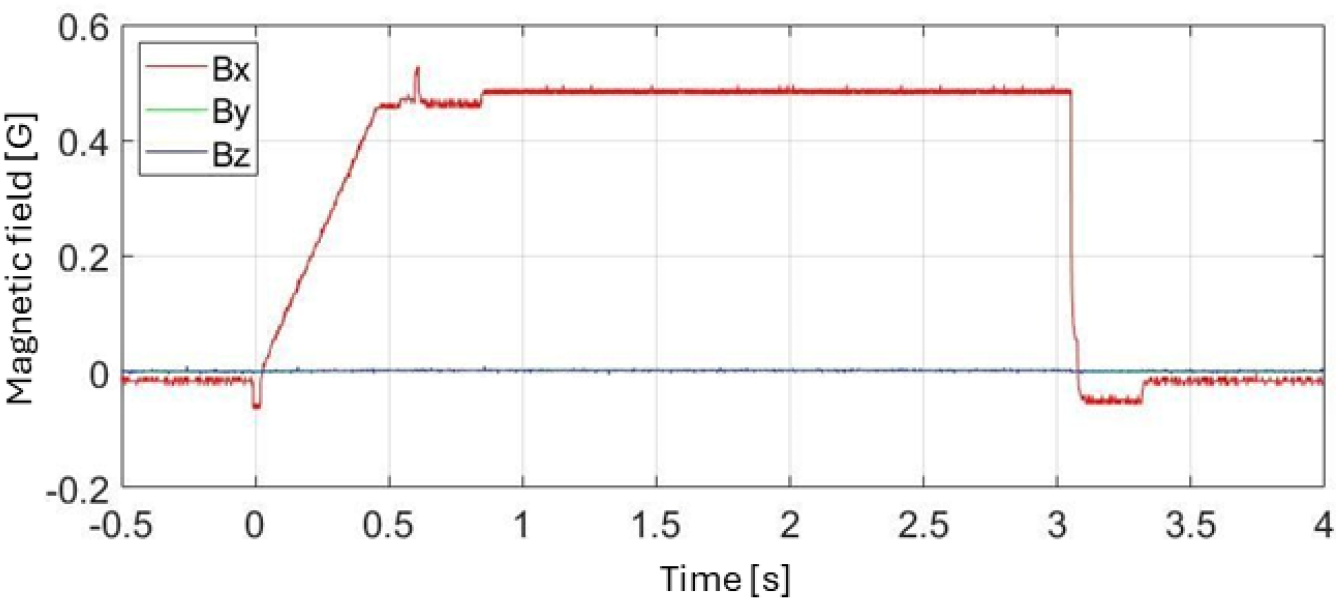
Magnetic field measurement along the *X*-axis during a 3-second 0.5 G magnetic field stimulus with a 0.5-second ramp-up time.

**Fig. 9.**
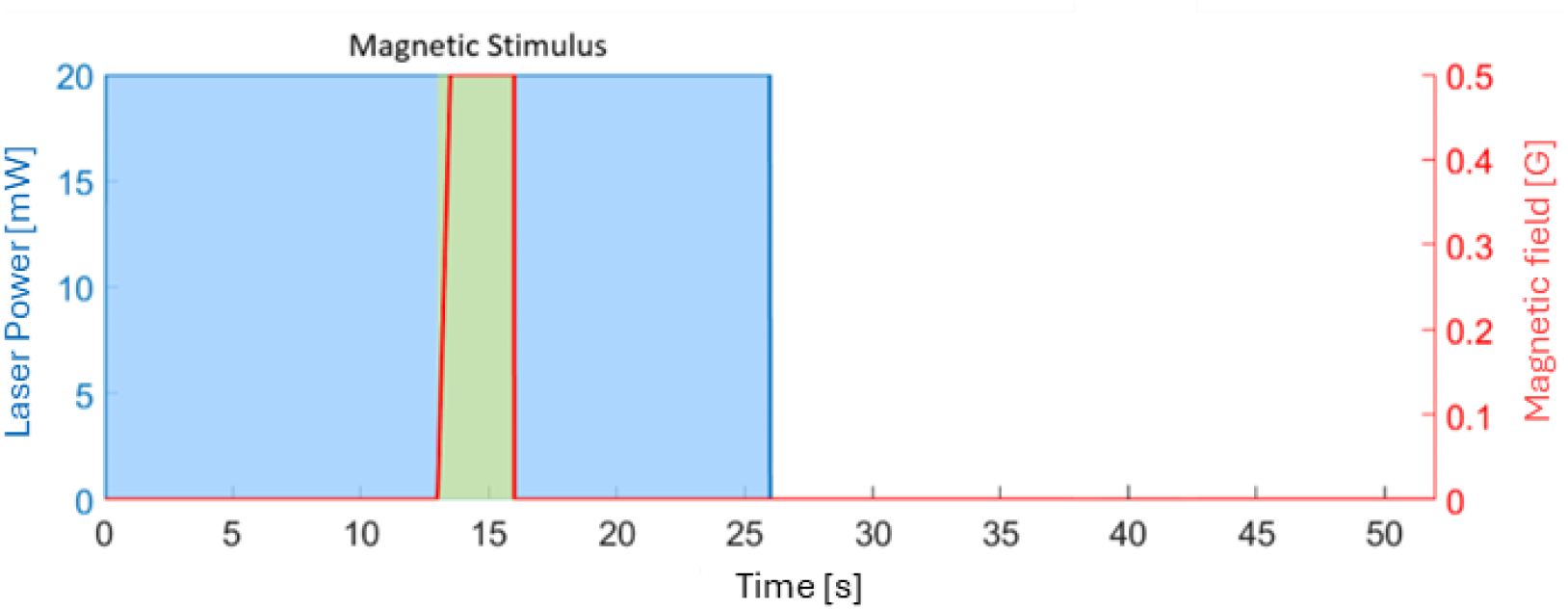
Pulse sequences for the magnetic response experiment, including visual (blue) and magnetic (red) stimuli. Magnetic field strength and blue laser power at the fiber output are indicated.

### 2.4 Calcium Imaging

#### 2.4.1 Microscopy setup

Calcium imaging was performed using a two-photon microscope (Ultima IV, Bruker) (Fig. 6) equipped with an ultra-short pulsed laser (Mai Tai DeepSee HP, Spectra-Physics). Excitation intensity was controlled by a Pockels cell, and scanning was performed with a pair of galvo mirrors. The microscope operated in epi-fluorescence configuration with a 20× water-immersion objective (NA 1.0, XLUMPLFLN20XW, Olympus). Emitted fluorescence was filtered with a 525±35 nm bandpass filter (Chroma) and detected by a photomultiplier tube (PH11706-40, Hamamatsu). The whole microscope is optically shielded from the environment by a light-tight enclosure.

#### 2.4.2 Imaging parameters

The following image parameters yielded the best results, balancing spatial and temporal resolution with signal-to-noise ratio and photobleaching.

1. Imaging wavelength: An excitation wavelength of 780 nm maximized Fura fluorescence.
2. Laser power: The excitation laser power ranged from 4 to 5 mW, providing an optimal signal-to-noise ratio while minimizing photodamage, which was determined via the fluorescence signal decay over the entire experimental sequence.
3. Field of view: Given the approximate 120 µm width of the AOTu, images were acquired at 128×128 pixels with a 2×2 µm pixel size.
4. Exposure time: An exposure time of 6.4 µs per pixel provided a good signal-to-noise ratio while enabling recording at a 6 Hz frame rate.

The microscope data collection was triggered by the stimulus control computer, allowing for precise synchronisation between stimuli and recording.

### 2.5 Data Analysis

Two-photon calcium imaging data were analyzed with custom MATLAB scripts [60,61], directly after every experimental sequence, performing the following processing steps. The corresponding code is available on https://github.com/NeurophysicsTrento/BeeMagnetoreception.

#### 2.5.1 Data analysis pipeline

1. Multi-frame TIFF files from sequential scans were combined into a single video for each experimental sequence, representing fluorescence intensity across space and time. The average intensity across the images over time was examined to evaluate the photobleaching and the need for compensation. It also allowed to identify where motion artifacts or other fluctuations were dominating the signal, to eliminate these trials from further analysis.
2. The entire video sequence was averaged over time for a single reference image in which Regions of interest (ROIs) were selected. To assist this procedure, a regional homogeneity (ReHo) analysis can be performed, which highlights boundaries between regions of correlated signal fluctuations.
3. The fluorescence signal was then averaged within each ROI, which substantially reduces the signal-to-noise ratio. Optional processing steps included detrending and high-pass filtering, both to compensate for the slow exponential decay due to photobleaching, and deconvolution with the Fura signal decay function to compensate for the delay due to unbinding of calcium from the detector molecule.
4. The relative fluorescence change was computed by normalizing the fluorescence during the stimulus with the average baseline during a pre-stimulus phase, usually lasting 5 s. Because neural activation decreases the Fura fluorescence intensity, traces were inverted so that activated neurons provided positive signal changes. The resulting measure, −Δ*F/F*, is proportional to the neuronal firing rate [62].
5. −Δ*F/F* was analyzed across the single trials, providing mean responses and standard deviations. ROIs were labeled as active when the mean fluorescence changes during the stimulus period exceeded 1.96 × the pre-stimulus standard deviation, which corresponds to a 95% confidence threshold, visualized with blue-to-red shading. A proper statistical analysis was performed using paired *t*-tests between pre-, stimulus-, and post-stimulus periods, with false discovery rate (FDR) compensation.

## 3 Results

### 3.1 Structural data

Mosaicking two images acquired with a 20× objective enables visualization of the dye diffusion pathway from the injection side near the left AOT along the ventral inter-tubercle tract (vlTT) to the contralateral AOTu (Fig. 10).

**Fig. 10.**
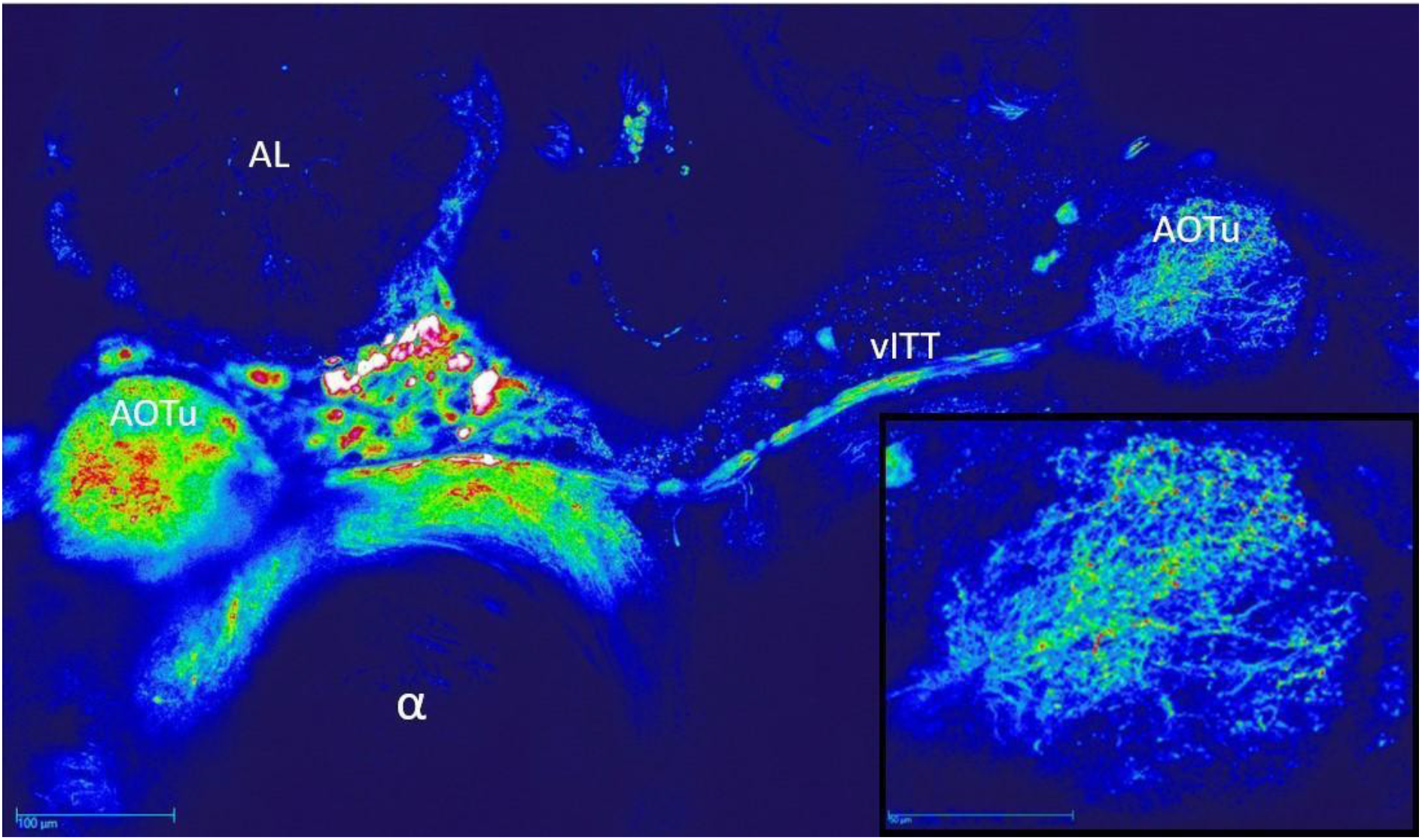
Image of the bee brain showing staining, with the bottom-right inset providing a zoomed view of the AOTu. The injection site is located in the left AOTu, with dye diffusing to the right AOTu via the vITT. The dimension of the full image is 425 × 240 μm2, in the inset 160 × 115 μm2. Abbreviations: AL: antennal lobe; α: mushroom body α-lobe; vITT: ventral inter-tubercle tract.

For each subject, the volumetric structure of the AOTu was determined by recording Z-stacks at higher resolution after the functional imaging sequence, using 512×512-pixel images and a slice distance of 2.5 μm (Fig. 11). These data enable 3D reconstruction using magic wand image segmentation in Amira (V5.4, Thermo Scientific) (Fig. 11B).

**Fig. 11.**
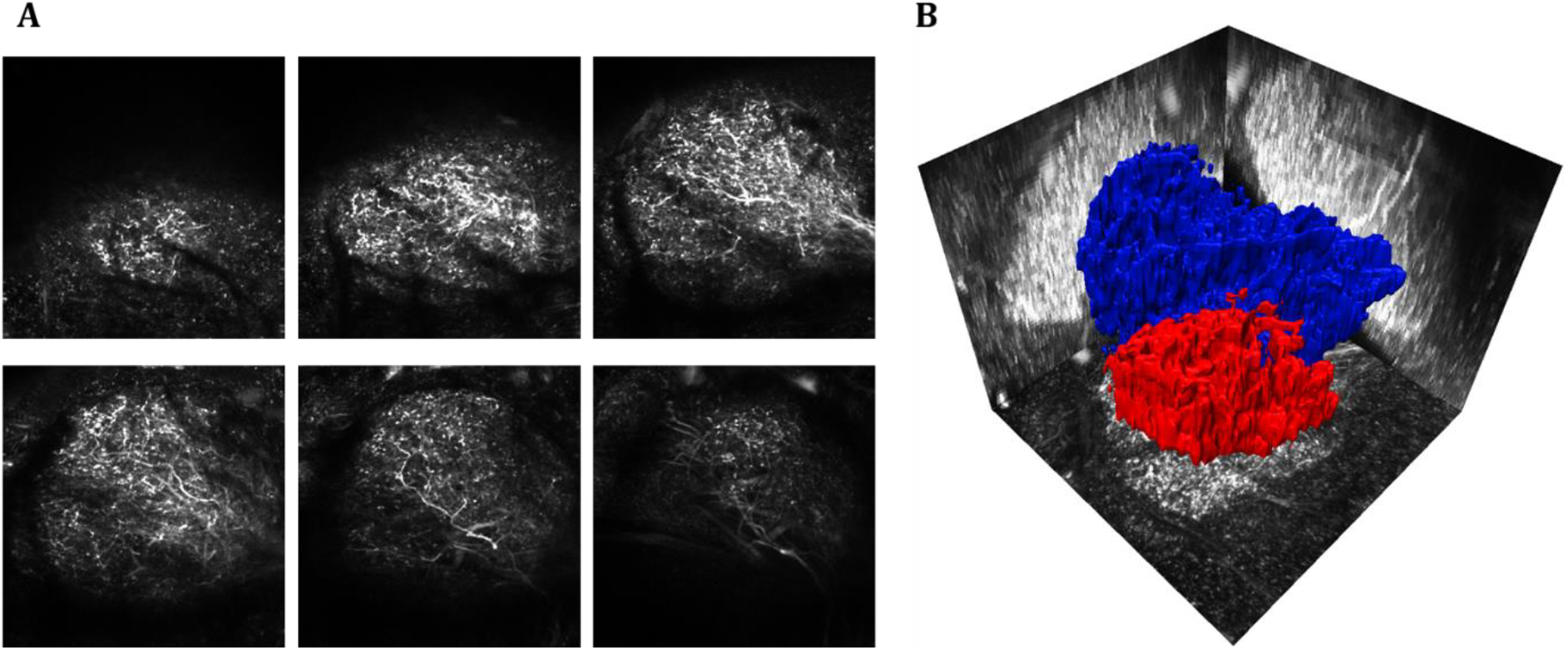
(A) Stack of images traversing the AOTu, image dimensions are 160 × 160 μm^2^, the distance between the shown images is 20 μm. (B) Reconstructed 3D structure of the same AOTu with the ventral lobe below and the dorsal lobe above, image dimensions are 160 × 160 × 140 μm^3^.

### 3.2 Light-sensitive responses

A typical AOTu response to the described visual stimulus is shown in Fig. 12A. It displays a temporal dynamics consistent with previous reports [56,63]. The calcium trace exhibits two peaks, an ON response at stimulus onset and an OFF response at the offset, as well as a lower tonic response during illumination. The percentage fluorescence changes for the ON and OFF responses reached 0.8% and 1.1%, respectively, with a tonic component of 0.5%, which is lower than previous widefield microscopy results (ON: 4%, OFF: 5%, tonic: 4%) [31]. This reduction is likely due to the thinner optical sectioning provided by two-photon microscopy.

**Fig. 12.**
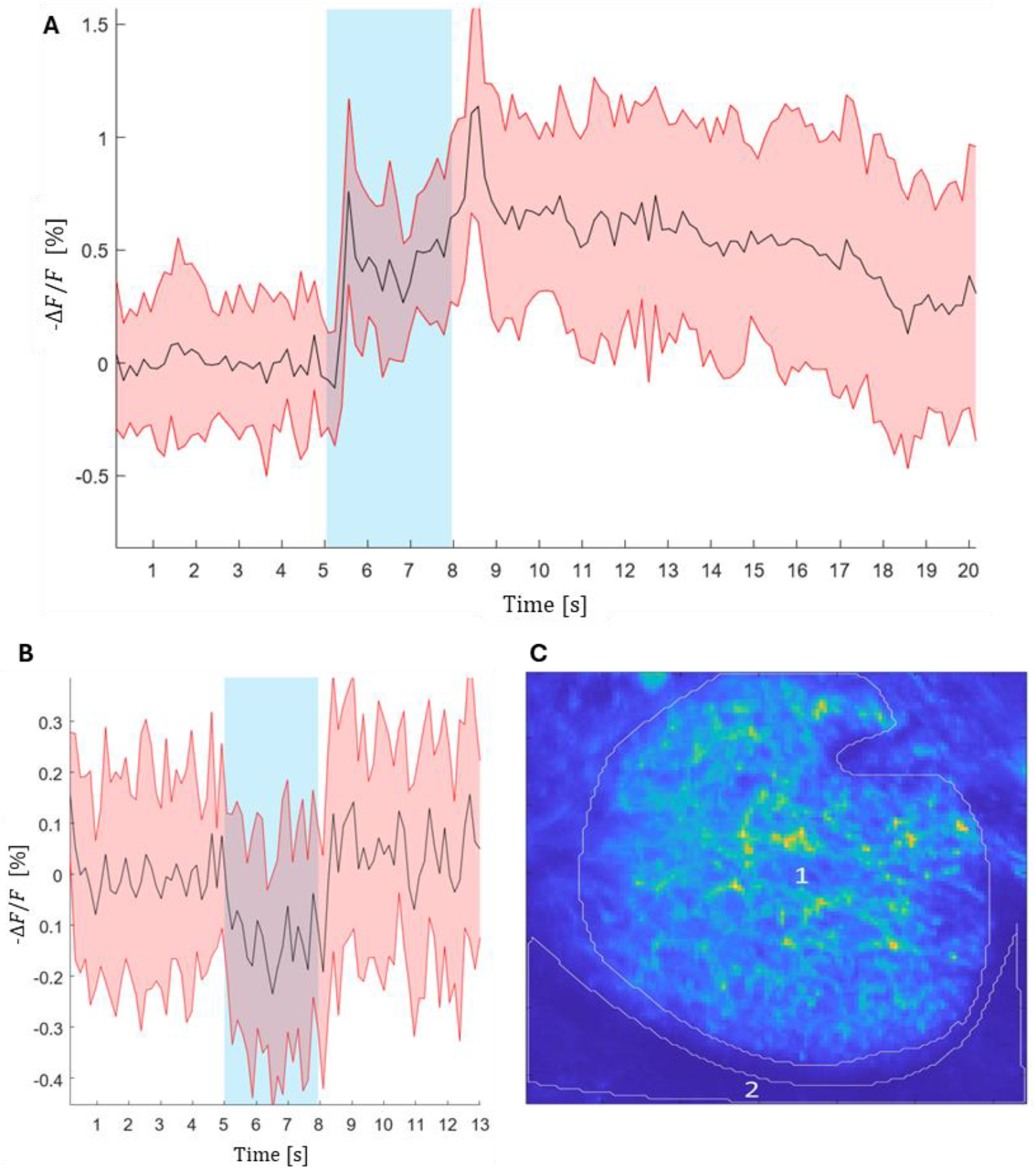
Time series of Fura fluorescence changes during the visual stimulus sequence, showing the percentage change in fluorescence averaged over 15 trials. (A) Fluorescence response of the AOTu to the stimulus (ROI1). (B) Fluorescence changes in the background region under the same stimulus (ROI2). (C) Fluorescence intensity image averaged across all frames, from which ROI1 and ROI2 were chosen. ROI1 showing the AOTu, where signal values were measured. ROI2 represents a reference area where absolute fluorescence is minimal, to determine the background signal from the visual stimulation light. Note that Fura fluorescence decreases in response to elevated neural activity. Fluorescence values are inverted for visualization, plotted as −Δ*F/F*, such that the background signal from the visual stimulus appears negative. Red curves indicate a significant change in fluorescence from the pre-stimulus baseline (95% confidence threshold).

Signal background changes due to the visual stimulus (Fig. 12B) were quantified in a region posterior to the AOTu (Fig. 12C, ROI 2) without significant fluorescence, revealing a 0.3% increase in −Δ*F/F* due to the laser light despite extensive mechanical and optical shielding. This background attenuated the apparent AOTu response amplitude, thus its precise value remains uncertain. Consequently, 0.3% was chosen as the upper limit for acceptable background change, constraining the maximum laser intensity used for visual stimulation.

### 3.3 Magneto-sensitive response

Fluorescence recordings from a bee subjected to the magnetic stimulation protocol after piecewise linear detrending are shown in Fig. 13A. Distinct visual ON and OFF responses remain clearly detectable. The tonic component was transiently suppressed after the on-response, and reemerged after approximately 2-3 s. At magnetic field onset, fluorescence initially decreased by about 0.2% within the first second and decayed with a similar time constant as the visual responses.

**Fig. 13.**
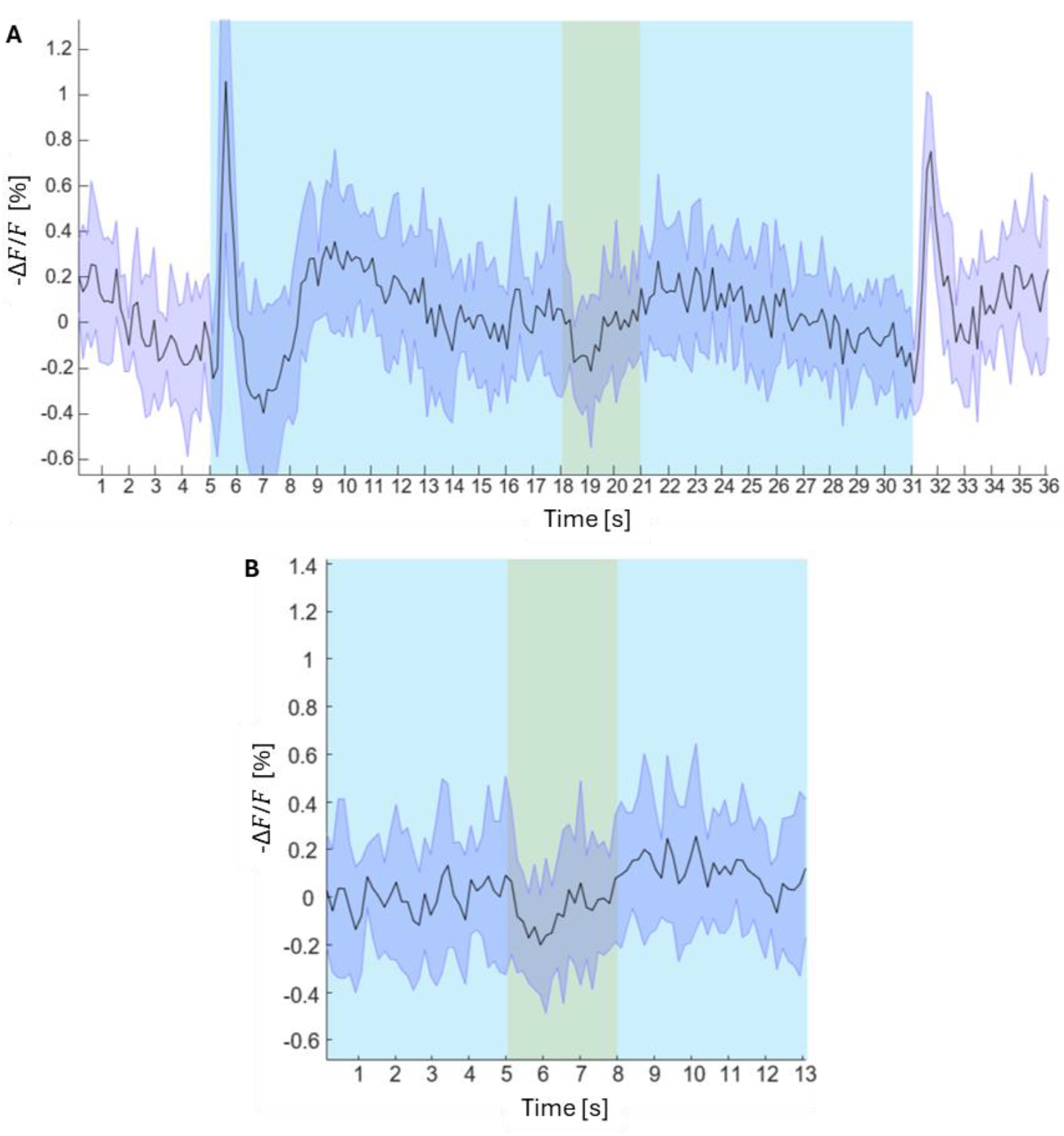
Detrended time-series recordings of calcium-sensitive fluorescence in the AOTu during magnetic stimulation, showing the percentage change in fluorescence −Δ*F/F* averaged over 10 trials. (A) Fluorescence response of the AOTu over the full stimulus sequence. (B) Response of the AOTu to the magnetic stimulus, in this recording, the fluorescence is normalized to the mean fluorescence during the pre-magnetic stimulus interval.

After the magnetic field offset, the signal returned to its pre-stimulus baseline. A paired *t*-test comparing pre-stimulus and stimulus fluorescence across trials (*n* = 10) yielded a *p*-value of 0.13, indicating that the average signal change during the entire stimulus period was not statistically significant.

## 4 Discussion

Several parameters were critical for obtaining reliable AOTu visual responses, including two-photon excitation power, visual stimulus intensity, and the imaging plane. The excitation power had to be carefully adjusted: While increasing the power amplified the fluorescence and reduced the relative background from the visual stimulus, it also amplified the noise and the photobleaching. Minor changes in the excitation laser power on the order of 0.2 mW were decisive for whether weak ON/OFF responses were visible, the signal was masked by noise (if the power was too high), or stimulus light artifacts dominated (if the power was too low). Using optimal laser intensities, peak AOTu activation signals of 1.2% were obtained. The imaging plane also affected response amplitude, with the most robust signals recorded 10 µm above the AOTu center.

Although parameters that yield pronounced ON and OFF responses were relatively straightforward to identify, obtaining a clear tonic component during stimulation proved substantially more challenging and also depended critically on overall preparation quality, being highly sensitive to both dye loading and tissue integrity. Because the AOTu lies close to the brain surface, it was particularly vulnerable to damage during tracheal removal, and even minor, visually undetected damage frequently abolished the tonic response. In such cases, otherwise well-stained tubercles often exhibited only ON/OFF responses or no detectable activity at all.

Dye concentration imposed an additional constraint: insufficient staining increased noise due to dominant background autofluorescence, whereas excessive staining often eliminated detectable activity. In heavily stained tubercles, reducing excitation power to compensate for elevated baseline fluorescence did not restore robust responses, and at most, a sharp ON response was observed. This loss of signal may reflect excessive intracellular buffering by the calcium indicator Fura, attenuating activity-dependent calcium transients and thereby suppressing detectable responses [64].

The injection strategy also proved important: using three injections with borosilicate glass needles, each containing a small amount of dye, was preferable to a single injection with the same total dye load, because the latter often caused leakage into adjacent regions, such as the mushroom bodies, and increased background fluorescence. In contrast, three small injections at the same location limited background staining while still providing sufficient labeling of the tubercle.

To reliably detect a response to magnetic stimulation, a strong and stable visual response was required, characterized in particular by a high tonic response with low noise. Bees that lacked sustained visual activity showed no detectable change during magnetic stimulation. When a fluorescence change during magnetic stimulation was observed, it consistently manifested as a decrease, suggesting that this type of magnetic stimulus may suppress neural activity. The larger the visual response, the better the detectability of this inhibitive change.

The origin of this potential inhibition in brain activity in response to the magnetic stimulus remains to be determined, but RPM models indeed suggest a bright/dark modulation of the blue-light photoreceptor activity that is projected via the optic lobe into the AOTu [40]. Whether the AOTu can also show excitation in response to magnetic stimuli of different orientations remains to be investigated.

In most bees with a magnetic response, the tonic response to the visual stimulus during the magnetic pulse sequence initially dropped back toward pre-stimulus levels after the ON response and recovered after 2-3 s (Fig. 13A). This reduction was never observed during the pure visual stimulus sequence (Fig. 12A), and its origin remains unclear. The only systematic difference between the two stimulus protocols is that the magnetic sequence begins in a zero-field environment, suggesting that the AOTu responds differently to pure optical stimuli under hypomagnetic field conditions, though further experiments are needed to confirm this. It is also important to assess whether the magnetic response is influenced by the visual stimulus protocol, specifically whether the visual stimulus remains on across trials or alternates on and off during each trial. So far, only the latter has been tested (Fig. 9) to confirm that the AOTu remained equally responsive to visual stimuli during each trial.

## 5 Conclusion and outlook

The weak and variable nature of magnetic-field-dependent responses, which has made controlled experimental investigation particularly challenging across model systems [65], has also limited our study to largely qualitative results. However, this two-photon imaging approach presented here suggests the feasibility of detecting weak, magnetic-field-dependent modulation of neural activity in the honeybee brain. By enabling *in vivo* measurements in a higher-order visual processing region, this framework provides access to downstream neural circuits in which magnetically elicited signals may be integrated and amplified. The observation of small fluorescence changes in the AOTu may provide a basis for probing the neural correlates of magnetoreception beyond primary sensory structures.

Several technical improvements are expected to further enhance sensitivity and interpretability. First, the blue-light background signal that currently limits the detection of small fluorescence changes will be eliminated by implementing a flyback stimulation protocol, in which the blue laser is applied only during the galvanometric scanner retrace, which will also allow for increasing light stimulus intensity [66]. Second, the present measurements are restricted to a single imaging plane, whereas magnetically modulated activity is likely distributed across the three-dimensional structure of the AOTu. Extending the approach to volumetric imaging will allow reconstruction of spatial activity patterns and may reveal localized or distributed response motifs.

While the present results provide only preliminary evidence of magnetosensitive responses, the setup will allow for further investigation of the underlying mechanism, in particular, a controlled testing of key predictions of the radical pair mechanism models, including the requirement for short-wavelength (blue) light, the expected sensitivity to magnetic field axis rather than polarity, and the disruption of responses by weak radiofrequency fields, which can be easily integrated into the setup. Systematic variation of these parameters will be essential to determine whether the observed signals are consistent with a radical-pair–based mechanism.

Finally, these first findings highlight the importance of systematically linking stimulus design to underlying biological hypotheses. In particular, the predominance of inhibitory responses under the present conditions complicates the interpretation of light dependence and suggests that stimulus configurations may need to be optimized to elicit excitatory activity. Identifying such conditions will be critical for disentangling the contributions of visual processing and magnetically induced modulation.

Overall, this imaging framework provides a new methodological approach for studying magnetoreception within intact neural circuits, bridging the gap between molecular models and systems-level function. By enabling controlled, *in vivo* imaging of weak magnetic effects in the brain, it may open new avenues for testing mechanistic hypotheses and for dissecting how magnetic and visual information are integrated in the neural circuitry underlying animal navigation.

## 6 Author contributions: CRediT

Alan Oesterle: Data curation, Formal analysis, Investigation, Methodology, Software, Validation, Visualization, Writing – original draft, review and editing Alara Kiriş: Methodology, Investigation, Writing - review and editing Albrecht Haase: Conceptualization, Funding acquisition, Methodology, Project administration, Resources, Software, Supervision, Validation, Visualization, Writing - original draft, review and editing

## Acknowledgements

We acknowledge help with the magnetic field simulation by Riccardo Cabassi. A.O. acknowledges financial support from the Q@TN joint laboratory.

